# Effects of upgrading acquisition-techniques and harmonization methods: A multi-modal MRI study with implications for longitudinal designs

**DOI:** 10.1101/2021.10.31.466635

**Authors:** Takashi Itahashi, Yuta Y. Aoki, Ayumu Yamashita, Takafumi Soda, Junya Fujino, Haruhisa Ohta, Ryuta Aoki, Motoaki Nakamura, Nobumasa Kato, Saori C. Tanaka, Daisuke Kokuryo, Ryu-ichiro Hashimoto

## Abstract

A downside of upgrading MRI acquisition sequences is the discontinuity of technological homogeneity of the MRI data. It hampers combining new and old datasets, especially in a longitudinal design. Characterizing upgrading effects on multiple brain parameters and examining the efficacy of harmonization methods are essential. This study investigated the upgrading effects on three structural parameters, including cortical thickness (CT), surface area (SA), cortical volume (CV), and resting-state functional connectivity (rs-FC) collected from 64 healthy volunteers. We used two evaluation metrics, Cohen’s *d* and classification accuracy, to quantify the effects. In classification analyses, we built classifiers for differentiating the protocols from brain parameters. We investigated the efficacy of three harmonization methods, including traveling subject (TS), TS-ComBat, and ComBat methods, and the sufficient number of participants for eliminating the effects on the evaluation metrics. Finally, we performed age prediction as an example to confirm that harmonization methods retained biological information. The results without harmonization methods revealed small to large mean Cohen’s *d* values on brain parameters (CT:0.85, SA:0.66, CV:0.68, and rs-FC:0.24) with better classification accuracy (>92% accuracy). With harmonization methods, Cohen’s *d* values approached zero. Classification performance reached the chance level with TS-based techniques when data from less than 26 participants were used for estimating the effects, while the Combat method required more participants. Furthermore, harmonization methods improved age prediction performance, except for the ComBat method. These results suggest that acquiring TS data is essential to preserve the continuity of MRI data.

## Introduction

Structural and functional MRI (sMRI and fMRI) are primary non-invasive neuroimaging techniques for investigating the neural basis of psychiatric and neurological disorders [Nogovitsyn et al., 2020; Stephane et al., 2019]. To examine neural bases, several neuroimaging-based markers have been developed so far [Gordon et al., 2018; Prigge et al., 2018; Sintini et al., 2020; Wallace et al., 2015]. Because age is one of the major factors for psychiatric and neurological disorders, studies with longitudinal design are prevailing [Huang et al., 2020; Okada et al., 2019]. In addition, as MRI acquisition is costly and time-consuming, some cross-sectional studies need to recruit participants in a wide range of periods. In these cases, upgrading the protocol during the study is inevitable.

Upgrading the acquisition protocol improves the data quality and accuracy. Despite such a positive aspect, a downside of upgrading is discontinuity of the technical homogeneity of MRI data, which hampers longitudinal and long-lasting cross-sectional studies. Thus, combining the data acquired with the prior protocol and the data with the latest protocol is a challenge to conduct studies on aging successfully.

Although integrating two sets of MRI data is essential, previous studies have mainly investigated the impacts of MR system upgrading on sMRI data. For instance, upgrading the MRI scanner from Siemens TimTrio to Prisma fit showed increased cortical thickness (CT) values in the frontal, temporal, and cingulate cortices [Plitman et al., 2021; Potvin et al., 2019]. Despite the extensive use of resting-state fMRI (R-fMRI), only a few studies have examined the effects of upgrading acquisition protocol (e.g., choice of multi-band factors) on fMRI data [Demetriou et al., 2018; Risk et al., 2018; Risk et al., 2021; Srirangarajan et al., 2021].

The harmonization method may be one of the practical approaches to mitigate such effects. Several harmonization methods have been proposed [Beer et al., 2020; Fortin et al., 2017; Fortin et al., 2018; Maikusa et al., 2021; Yamashita et al., 2019]. These harmonization methods might be plausible to resolve discontinuities with existing MRI data, yet no prior studies have investigated their efficacy on MRI data before and after upgrading acquisition techniques. In addition, these methods require MRI data acquired from both protocols to estimate the measurement bias, and thus it is crucial to identify the minimum number of participants to avoid unnecessary costs and time.

The current study investigated the effects of upgrading acquisition protocols on structural parameters and resting-state functional connectivity (rs-FC) data collected from healthy adult volunteers. First, we performed univariate analyses to quantify differences of brain parameters using Cohen’s *d* and paired t-tests. We also applied classification analyses to ask whether brain parameters could provide information about protocols in a multivariate manner. We then used the three most frequently utilized harmonization methods, including TS [Yamashita et al., 2019], TS-ComBat [Maikusa et al., 2021], and ComBat [Fortin et al., 2017; Fortin et al., 2018; Yu et al., 2018], to investigate whether these methods alleviate the effects of protocol upgrades and how many participants are necessary for estimating the impacts. We selected these methods because of their simplicity and applicability. Finally, we performed age prediction to investigate whether harmonization methods preserve critical biological information.

## Materials and Methods

### Participants

Sixty-five healthy adult volunteers [2 females; mean age ± standard deviation (SD): 32.3 ± 7.5 years old); age range: 20–45 years] participated in this study. Participants underwent two MRI sessions using a 3T MRI scanner (MAGNETOM Verio dot, Siemens Medical Systems, Erlangen, Germany). In one session, we used a single-band fMRI acquisition protocol used in a multi-site, multi-disease cohort study [Strategic Research Program for Brain Science (SRPBS) protocol] with a 12-ch head coil [Tanaka et al., 2021]. We used a multiband fMRI acquisition (Harmonized Protocol; HARP) with a 32-ch head coil in the other session. The HARP protocol has been developed for minimizing the scanner differences [Koike et al., 2021]. Of note, 30 out of 65 participants underwent an MRI session with the HARP protocol within one month after an MRI session with the SRPBS protocol because the HARP protocol was not available at that time. We excluded one male participant from our analyses because of excessive head motion during the scans. We provided the MRI acquisition parameters in Table S1.

The study was approved by the Institutional Review Board of Showa University Karasuyama Hospital and was prepared in accordance with the ethical standards of the Declaration of Helsinki. Written informed consent was obtained from all the participants after fully explaining the purpose of this study.

### Structural MRI preprocessing

We preprocessed sMRI data using FreeSurfer version 6.0.1. The details of preprocessing steps are described in detail in earlier studies [Dale et al., 1999; Fischl et al., 1999]. Briefly, this software performed a series of preprocessing procedures, including spatial normalization, bias field correction, intensity normalization, skull-stripping, segmentation, and reconstruction of surface mesh. For each participant, we then computed three parameters: cortical thickness (CT), surface area (SA), and cortical volume (CV). To characterize the structural characteristics, we used Schaefer’s 400 cortical parcels [Schaefer et al., 2018] as regions of interest (ROIs) to extract the mean values for the three parameters. These procedures yielded a 400-dimensional feature vector for each structural parameter in each participant.

### R-fMRI data preprocessing

We preprocessed R-fMRI data using FMRIPREP version 1.1.8 [Esteban et al., 2019]. We used the same preprocessing pipeline to avoid bias introduced by differences in the pipeline. FMRIPREP performs a series of preprocessing steps, including head motion estimation, slice timing correction, co-registration of echo-planar image data to the corresponding T1-weighted anatomical image, distortion correction, and normalization to a standard Montreal Neurological Institute (MNI) space. To analyze the preprocessed data using the Human Connectome Project (HCP) style surface-based methods, we used the ciftify toolbox version 2.1.1 [Dickie et al., 2019] that allowed us to analyze our non-HCP style data using an HCP-like surface-based pipeline.

For each vertex, we performed nuisance regression to remove the effects of artifactual and non-neural sources. Nuisance regressors consisted of six head-motion parameters, averaged signals from subject-specific white matter and cerebrospinal fluid masks, global signal, their temporal derivatives, and linear detrending. After nuisance regression, we applied a band-pass filter (0.008–0.08 Hz) to the residuals. We computed framewise displacement (FD) [Power et al., 2012] for each participant to characterize the frame-by-frame head motion during the scans. We used FD as a measure for detecting occasional head movement. To reduce spurious changes in FC due to head motion, we removed volumes with FD > 0.5 mm, as proposed in a previous study [Power et al., 2012].

We used Schaefer’s 400 cortical atlas [Schaefer et al., 2018] in combination with 17 subcortical regions [Fischl et al., 1999] and 10 cerebellar regions [King et al., 2019] as ROIs to characterize the whole-brain connectivity pattern. The Pearson correlation coefficient was computed among all possible pairs of ROIs to characterize the functional connectome, resulting in a 427 × 427 functional connectivity matrix for each participant.

### Evaluation measures

We computed three measures to characterize the effects of protocol upgrades and assess how the harmonization methods mitigate the effects.

#### Cohen’s *d*

We computed the effect size (Cohen’s *d*) to characterize the effects of protocol upgrades on brain parameters. The Cohen’s *d* for the *j*-th brain parameter, *d_j_*, is computed as

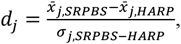

where *σ_SRPBS–HARP_* stands for the standard deviation of the difference from the SRPBS protocol to the HARP one; brain parameter with positive Cohen’s *d* value indicates that the brain parameter of the SRPBS protocol shows a higher value than that of the HARP protocol.

#### Classification accuracy

To investigate how the classification accuracy changes before and after applying the harmonization methods, we performed classification analyses using three machine learning algorithms: logistic regression with the least absolute shrinkage and selection operator (LASSO) [Tibshirani, 1996], a ridge logistic regression, and a support vector machine (SVM). We used a 10-fold nested cross-validation scheme similar to our previous study [Yamagata et al., 2019]. We divided participants into ten folds in the 10-fold cross-validation framework while keeping the pair information. Of note, the term “pair” refers to the same subject in different datasets (i.e., SRPBS and HARP). We used all-but-one folds as training data to train classifiers in each fold, while we treated the remaining fold as test data for testing the classification performance. We then evaluated the impacts of protocol differences using classification accuracy.

We used the “*lassoglm*” function implemented in MATLAB (2020b, Mathworks, USA) for the LASSO method. In this function, we set “NumLambda” to 25 and “CV” to 10. In the inner loop, this function first computes a value of λ that is just large enough such that the only optimal solution is an all-zero vector. This function then creates a total of 25 equally spaced λ values from 0 to λ_max_. It then determines the optimal λ according to the one-standard-error rule. This function selects the largest λ within the standard deviation of minimum prediction error among all λ. For ridge logistic regression, we used the “*fitclinear*” function implemented in MATLAB. We used the “*fitcsvm*” function implemented in MATLAB for the SVM classifiers. We set “KernelFunction” to ‘linear’ and “OptimizeHyperparameters” as ‘BoxConstraint’ and ‘KearnelScale.’

#### Prediction performance

We performed age prediction as an example to investigate whether the harmonization methods retained age information. We used two machine learning algorithms for these analyses: support vector regression (SVR) and linear regression with the LASSO method. Of note, we did not apply ridge regression here because of poor prediction performance. Similar to the classification analyses, we used 10-fold nested cross-validation while preserving the pair information. In each fold, we used all-but-one folds as training data to construct a prediction model, while we treated the remaining fold as test data for testing the prediction performance.

We then computed the Pearson correlation coefficient between predicted age and actual age to evaluate the prediction performance. Once the prediction results were obtained from the data before and after harmonization methods, we computed the percentage of improvement on the prediction performance.

We used the “*lassoglm*” and “*fitrsvm*” functions implemented in MATLAB (2020b, Mathworks, USA) for the LASSO and the SVR, respectively. We set the hyperparameters similar to those used in the classification analyses.

### Harmonization methods

We used three harmonization methods to remove the protocol bias from brain parameters: TS, ComBat, and TS-ComBat methods.

#### Traveling subject (TS) harmonization method

The TS method is an extension of the general linear model (GLM) based method for correcting the protocol bias [Yamashita et al., 2019]. The uniqueness of this method is to use TS data. We estimated the participant factor and measurement bias (i.e., protocol bias) by fitting a linear regression to each brain parameter. We used a 1-of-*K* binary coding scheme for the participant factor and protocol bias. Let us consider the *<u>i</u>*-th participant from the *m*-th protocol. The corresponding coding vector, ***x**_im_*, becomes a row vector whose the *m*-th element is one and the rest are zeros. Similarly, the coding vector for the participant factor, ***x**_ip_*, is a row vector whose the *p*-th element is one and the rest are zeros. For each brain parameter, we considered the following regression model:

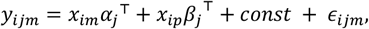

where *y_ijm_* represents the *j*-th feature for the *i*-th subject with the *m*-th protocol. The vectors, *α* and *β*, represent the protocol bias and participant factor, respectively. The superscript, ⊤, stands for the transpose of a vector or matrix.

We estimated the measurement bias and participant factors under the constraints such that the mean values of the participant factor and measurement bias are zero. We used the “*quadprong*” function implemented in MATLAB (R2020b, Mathworks, USA) for estimation. In contrast to the original TS method [Yamashita et al., 2019], the regression model did not incorporate sampling bias inside the design matrix. Thus, we did not add any regularization. After estimating the protocol bias and participant factor are computed, the harmonized feature, 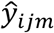, is calculated by subtracting the protocol bias from the brain parameters such that

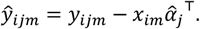

#### ComBat harmonization method

The ComBat method is originally proposed for correcting the batch effects in microarray data [Johnson et al., 2007]. This method has been used for adjusting the site effects in neuroimaging data [Fortin et al., 2017; Fortin et al., 2018; Yu et al., 2018] because of its simplicity and effectiveness. The ComBat method is based on location and scale adjustment model:

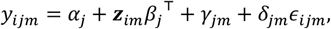

where *α_j_* is the overall constant term for the *j*-th feature, ***z**_im_* is the row vector whose elements are covariates of interest (e.g., age, sex, and disease status), and *β_j_* is a feature-specific vector of regression coefficients corresponding to ***z**_im_*. The terms, *γ_jm_* and *δ_jm_* represent the additive and multiplicative protocol effects of the *m*-th protocol for the *j*-th feature, respectively. In the current study, we incorporated age and sex as covariates of interest into the ComBat model. The ComBat method uses an empirical Bayes framework to estimate the bias terms, *γ_jm_* and *δ_jm_*. Finally, the *j*-th harmonized feature for the *i*-th participant from the *m*-th protocol is computed as

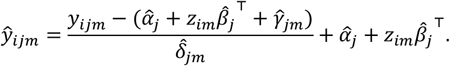

#### TS-ComBat harmonization method

The TS-Combat method is a recently-developed method for adjusting the site effects using TS data in the framework of the ComBat method [Maikusa et al., 2021]. The TS-Combat method replaces the row vector, ***z**_im_*, with the coding vector for the participant factor, ***x**_ip_*, such that

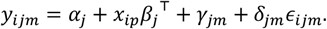

In contrast to the TS method, the TS-ComBat method does not incorporate any constraints on the participant factor. This method uses the Moore-Penrose pseudo inverse matrix to avoid the problem of rank deficiency in the design matrix. After estimating the coefficients, the *j*-th harmonized feature for the *i*-th participant from the *m*-th protocol is computed as

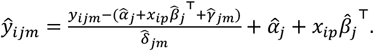

### Effects of the number of participants used for the estimation of protocol bias

We conducted additional analyses to investigate how many participants are necessary to estimate the effects of protocol bias. First, we randomly selected a subset of participants and fitted a GLM to assess the impacts of protocol upgrades. We repeated this random selection ten times. After subtracting the protocol bias from each brain parameter, we computed Cohen’s *d* and performed classification analyses as functions of the proportion of participants used in the harmonization method. The ratio of participants used to estimate the protocol bias was varied from 10% to 100% in 10% increments.

## Results

### Effects of protocol upgrades on Cohen’s *d*

#### Structural parameters

Figures 1A and 1C showed that the HARP protocol exhibited increased CT and CV values (i.e., negative Cohen’s *d* values) in the frontal regions and insular cortices compared to the SRPBS protocol. In contrast, the SRPBS protocol exhibited increased SA values (i.e., positive Cohen’s *d* values) in the frontal pole and orbitofrontal cortex (OFC) compared to the HARP protocol (Figure 1B). Before applying the harmonization methods, we observed medium to large mean Cohen’s *d* values (CT: 0.85 ± .23, SA: 0.66 ± 0.19, and CV: 0.68 ± 0.17 [mean ± SD]).

**Figure 1.**
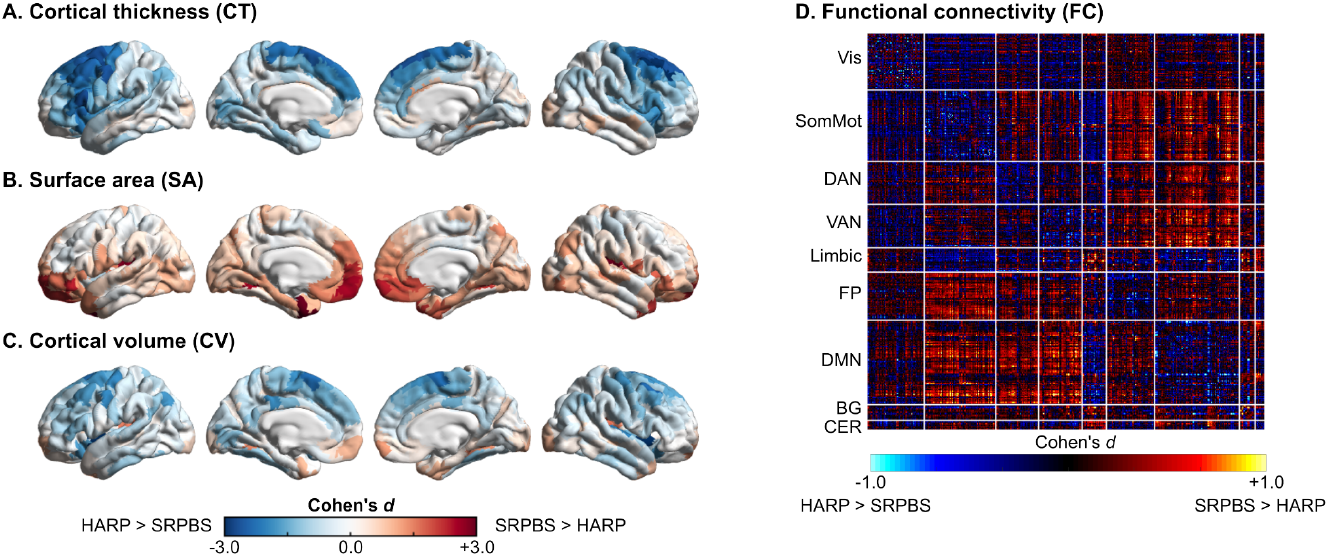
The upgrading effects on brain parameters. (A) cortical thickness (CT), (B) surface area (SA), (C) cortical volume (CV), (D) and functional connectivity (FC). The red color indicated that the SRPBS protocol exhibited significantly higher values in brain parameters compared to the HARP protocol, while the blue color indicated that the HARP protocol exhibited significantly higher values in brain parameters compared to the SRPBS protocol. Cohen’s *d* was computed in each brain region and functional connectivity. **Abbreviations:** BG: basal ganglia, CER: cerebellar, DAN: dorsal attention network, DMN: default mode network, FP: fronto-parietal network, SomMot: somatomotor network, VAN: ventral attention network, and Vis: visual network.

#### rs-FCs

Figure 1D showed that, compared to the HARP protocol, the SRPBS protocol showed higher rs-FC strengths (positive Cohen’s *d* values) stemming from default mode network (DMN) to other networks (e.g., somatomotor, dorsal attention [DAN], and ventral attention networks [VAN]). In addition, the SRPBS protocol showed decreased rs-FC strength (negative Cohen’s *d* values) in within-network connections, except for the limbic network. Before applying the harmonization methods, we observed a small mean Cohen’s *d* value (0.24 ± 0.06 [mean ± SD]).

### Effects of protocol upgrades on classification accuracy

#### Structural parameters

As shown in Table 1, the three classifiers achieved higher classification performance (classification accuracy > 92%) for all the structural parameters before applying the harmonization methods.

**Table 1.** Classification accuracy.

| Accuracy [%] | SVM | LASSO | Ridge |
| --- | --- | --- | --- |
| CT | 98.44 | 100 | 100 |
| SA | 93.75 | 98.44 | 94.53 |
| CV | 93.75 | 96.88 | 92.19 |
| rs-FC | 98.44 | 99.22 | 96.88 |
| **Abbreviations:** CT: cortical thickness, CV: cortical volume, SVM: support vector machine, rs-FC: resting-state functional connection, |  |  |  |

#### rs-FCs

Similar to the results of structural parameters, all the classifiers exhibited higher classification performance (> 96% accuracy) (Table 1).

### Effects of protocol upgrades on age prediction

#### Structural parameters

Before applying the harmonization methods, prediction models showed the following prediction performance (CT: *r*_LASSO_ = 0.27 and *r*_SVR_ = 0.33; SA: *r*_LASSO_ = 0.49 and *r*_SVR_ = 0.47; and CV: *r*_LASSO_ = 0.61 and *r*_SVR_ = 0.54) (Table 2).

**Table 2.**
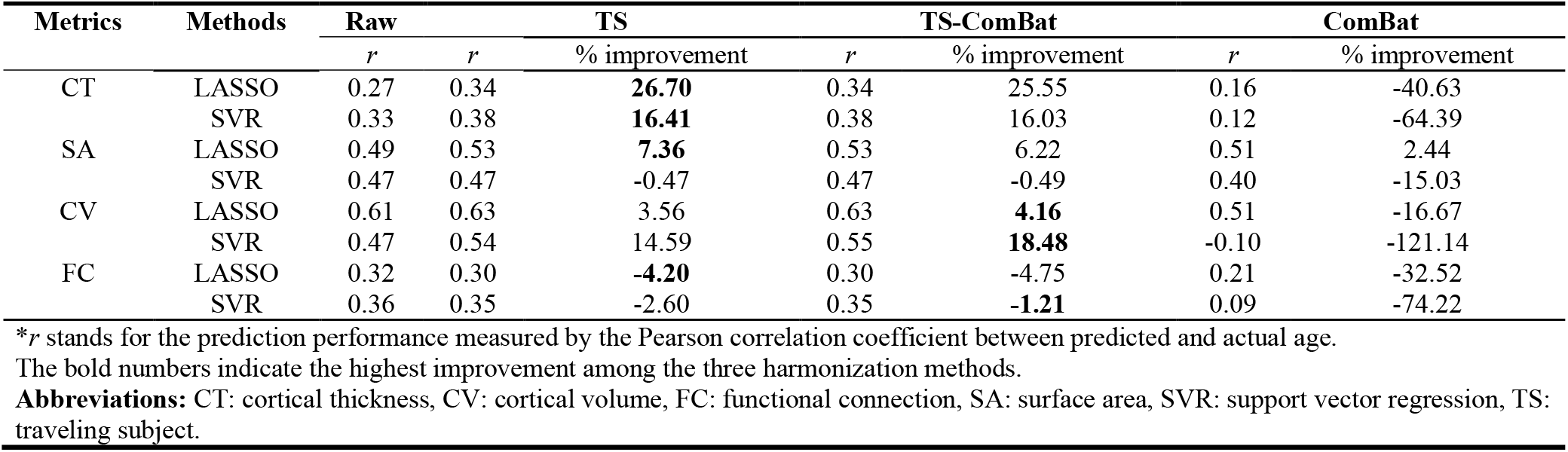
Effects of harmonization methods on age prediction.

#### rs-FCs

As shown in Table 2, prediction models showed the following performance (*r*_LASSO_ = 0.32 and *r*_SVR_ = 0.36) before applying the harmonization methods.

### Effects of the harmonization methods on Cohen’s *d*

#### Structural parameters

As shown in Figures 2A-C, all the harmonization methods reduced Cohen’s *d* values of all the structural parameters. In the TS and TS-ComBat methods, Cohen’s *d* values approached zero for all the structural parameters when increasing the number of participants used for estimating the protocol bias. In the ComBat method, Cohen’s *d* values could not reach zero (CT: 0.19 ± 0.05, SA: 0.19 ± 0.05, and CV: 0.32 ± 0.08).

**Figure 2.**
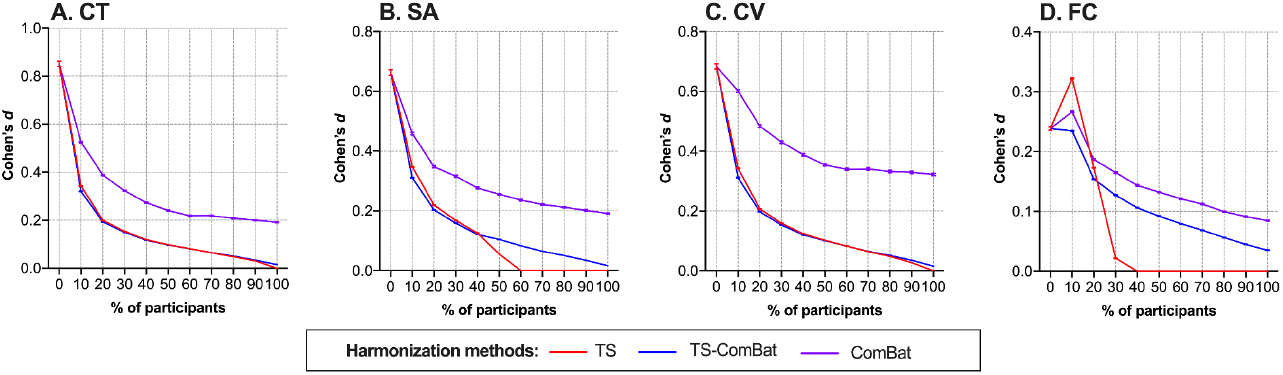
The effects of three harmonization methods on brain parameters measured by Cohen’s *d*. The standardized effect size was computed using Cohen’s *d* for four brain parameters: (A) cortical thickness (CT), (B) surface area (SA), (C) cortical volume (CV), and (D) functional connectivity (FC). We varied the number of participants used for estimating the protocol bias from 0% to 100%. Of note, “0%” indicates that no harmonization methods are applied. The error bars indicated the standard error of the mean (SEM). **Abbreviations:** TS: traveling subject.

#### rs-FCs

Cohen’s *d* values were decreased when applying harmonization methods. By applying the TS method, Cohen’s *d* values reached zero when 40% of participants were used (Figure 2D). In contrast, Cohen’s *d* values could not reach zero (TS-ComBat: 0.03 ± 0.01; and ComBat: 0.08 ± 0.02)

### Effects of harmonization methods on classification accuracy

#### Structural parameters

For all the structural parameters with TS and TS-ComBat methods, the classification performance of the LASSO method reached the chance level (i.e., 50%) when 20% to 30% of participants were used for estimating the protocol bias. In contrast, the LASSO method with the ComBat method required more participants to reach the chance level (the upper panels in Figure 3). Although the classification performance of SVM classifiers with three harmonization methods decreased in all the structural parameters, those for CT with TS and TS-ComBat methods decreased below 50% when more than half of the participants were used for estimating the protocol bias (the middle panels in Figure 3). The classification performance decreased below 50% for ridge logistic regression when more than 50% of participants were used (the lower panels in Figure 3).

**Figure 3.**
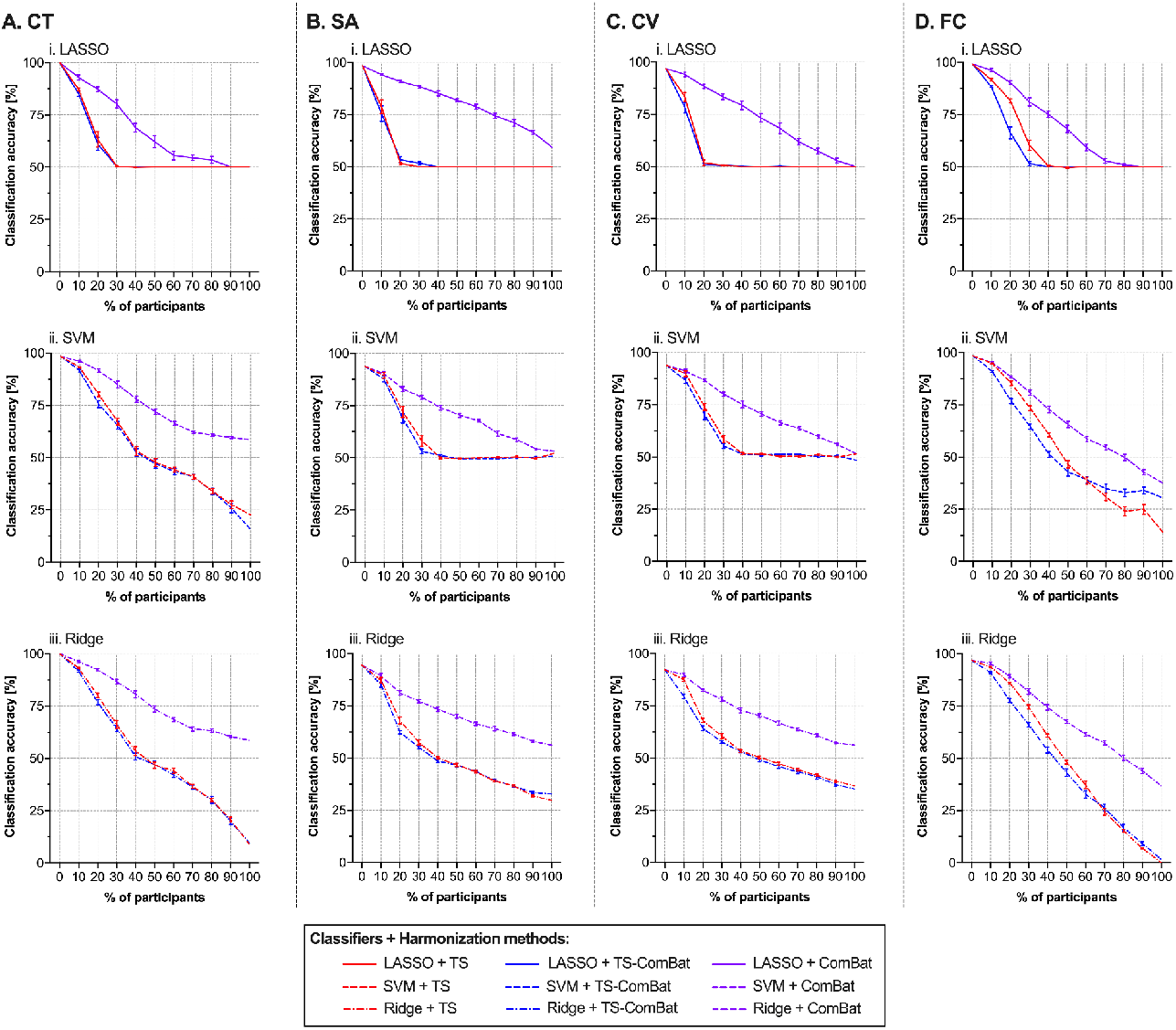
The effects of the harmonization methods on the classification analyses. We evaluated the classification performance of three classifiers on four brain parameters: (A) cortical thickness (CT), (B) surface area (SA), (C) cortical volume (CV), and (D) functional connectivity (FC). We varied the number of participants used for estimating the protocol bias from 0% to 100%. Of note, “0%” indicates that no harmonization methods are applied. The upper panels show the results of logistic regression with LASSO; the middle panels show the results of support vector machine (SVM); the lower panels show those of ridge logistic regression. We used classification accuracy as an index for classification performance. The error bars indicated the standard error of the mean (SEM).

We showed the distribution of posterior probabilities for ridge logistic regression with CT after applying the TS and TS-ComBat methods as an example (see Figure S1). In these results, we used all the participants to estimate the protocol bias. The posterior probabilities for both protocols were distributed around 0.5 with opposite directions, indicating a possibility of information leakage.

#### rs-FCs

Similar to the structural parameters, the classification of the LASSO method reached the chance level when 30% to 40% of participants were used for estimating the protocol bias (Figure 3D).

In contrast to the TS and TS-ComBat methods, the ComBat method required 90% of participants to reach the chance level. Similarly, the classification performance of SVM and ridge logistic regression decreased below the chance level.

We also showed the distribution of posterior probabilities for ridge logistic regression with FC after applying the TS and TS-ComBat methods (see Figure S2). In these results, the protocol biases were estimated using all the participants. The posterior probabilities for both protocols were distributed around 0.5 with opposite directions, indicating a possibility of information leakage.

### Effects of harmonization methods on age prediction

#### Structural parameters

Table 2 and Figures 4A–4C show the results of age prediction performance. After applying the harmonization methods, the LASSO methods exhibited improved prediction performance for all the structural parameters (*r*_CT_ = 0.34, *r*_SA_= 0.53, and *r*_CV_ = 0.63 for the TS and the TS-ComBat methods), except for the ComBat method (*r*_CT_ = 0.16, *r*_SA_= 0.51, and *r*_CV_ = 0.51). By applying the TS and TS-ComBat methods, SVR showed improved prediction performance for the CT and CV (TS: *r*_CT_ = 0.38, and *r*_CV_ = 0.54; TS-ComBat: *r*_CT_ = 0.38, and *r*_CV_ = 0.55), but not for SA (TS: *r*_SA_= 0.47; and TS-ComBat: *r*_SA_= 0.47). For the ComBat method, SVR showed decreased prediction performance in all the structural parameters (*r*_CT_ = 0.12, *r*_SA_= 0.40, and *r*_CV_ = −0.10).

**Figure 4.**
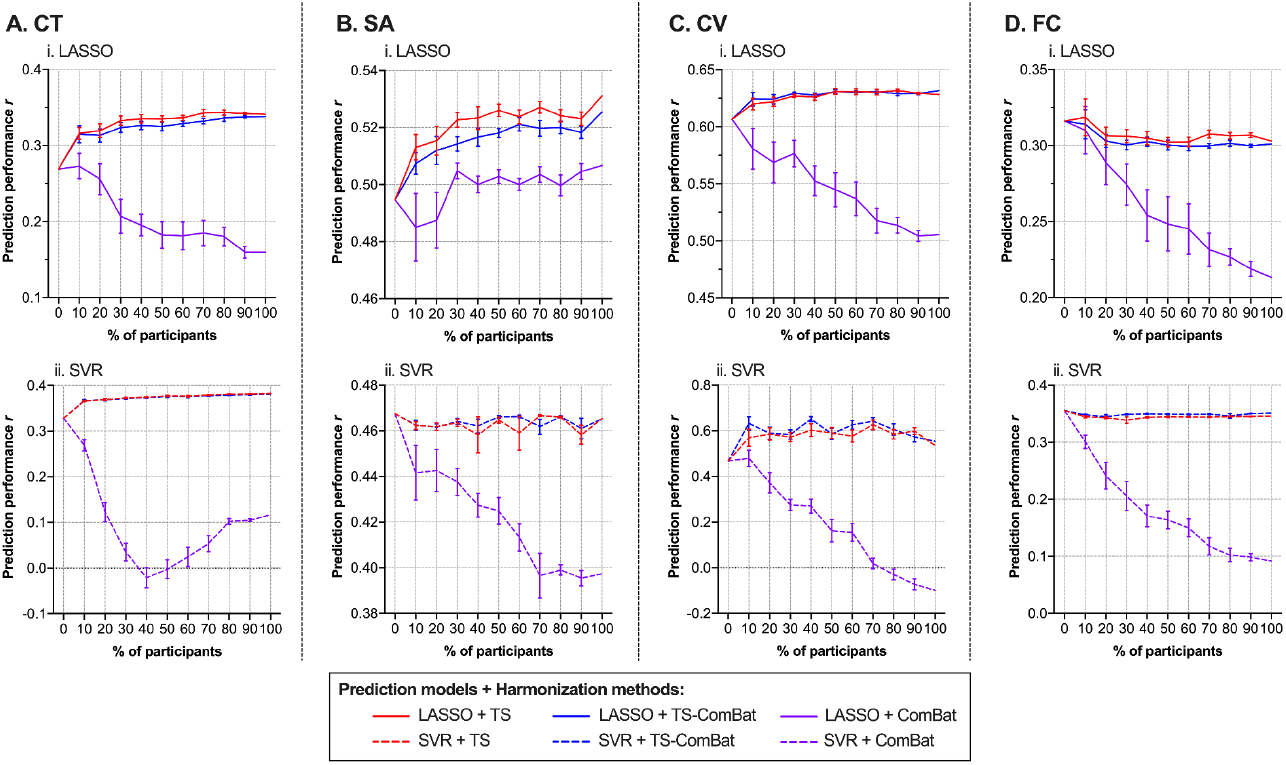
The effects of the harmonization methods on the age prediction. We evaluated the prediction performance of two prediction models on four brain parameters: A) cortical thickness (CT), (B) surface area (SA), (C) cortical volume (CV), and (D) functional connectivity (FC). We varied the number of participants used for estimating the protocol bias from 0% to 100%. Of note, “0%” indicates that no harmonization methods are applied. The upper panels show the results of logistic regression with LASSO, and the lower panels show the results of support vector regression (SVR). We used the Pearson correlation coefficient between predicted and actual age as an index for prediction performance. accuracy as an index for classification performance. The error bars indicated the standard error of the mean (SEM).

#### rs-FCs

In contrast to the structural parameters, the prediction performance were deteriorated by applying the harmonization methods (TS: *r*_LASSO_ = 0.30 and *r*_SVR_ = 0.35; TS-ComBat: *r*_LASSO_ = 0.30 and *r*_SVR_ = 0.35; and ComBat: *r*_LASSO_ = 0.21 and *r*_SVR_ = 0.09) (Figure 4D).

## Discussion

This study investigated the effects of upgrading acquisition techniques and harmonization methods on structural parameters and rs-FCs. Before applying the harmonization methods, we showed the impacts of protocol upgrades by Cohen’s *d* values and the classification accuracies. We also observed reduced upgrading effects by using three harmonization methods (i.e., TS, TS-ComBat, and ComBat). The TS and TS-ComBat methods showed that classification accuracy dropped to the chance level if data from 19 to 26 participants were available. On the other hand, the ComBat method required more participants to achieve the same level of performance. Furthermore, except for the Combat method, the harmonization methods improved the performance of age prediction using the structural parameters. In contrast, prediction models with rs-FCs could not improve the prediction performance after applying harmonization methods. These results suggest that the harmonization methods are promising methods for resolving the discontinuities with existing MRI data, especially sMRI, and TS data from 19 to 26 participants might be necessary before upgrading acquisition techniques.

Prior studies showed systematic effects of MRI scanner upgrades on the structural parameters, especially CT [Medawar et al., Plitman et al., 2021; Potvin et al., 2019]. The HARP protocol exhibited higher CT values in the medial, superior, and middle frontal gyri bilaterally, compared with the SRPBS protocol, those of which are in line with prior findings on upgrading MRI scanners. To complement our results on Cohen’s *d*, we assessed the effects of upgrading the acquisition techniques using the intra-class correlation (ICC) coefficients (see Supplementary Information). Compared with other structural parameters, CT showed relatively poor ICC coefficients (mean ICC = 0.49 for CT; Figure S3), which is consistent with prior findings [Iscan et al., 2015; Potvin et al., 2019]. FreeSurfer computes a CT value as a distance metric between white matter and pial surfaces [Fischl and Dale, 2000]. This metric, thus, is sensitive to the image quality (e.g., contrast-to-noise ratio). The improved image quality due to the protocol upgrades may offer the benefits of accurate estimation of these surfaces.

The current study observed a small mean Cohen’s *d* value in rs-FCs compared with the structural parameters. The HARP protocol exhibited lower rs-FC values in the basal ganglia (BG) and limbic networks, those networks of which might be sensitive to the scanner effects [Yamashita et al., 2019] and multiband acceleration [Risk et al., 2021]. This metric also showed poor ICC coefficients (mean ICC = 0.31; Figure S4), which is consistent with previous studies [Noble et al., 2019; Wang et al., 2017]. Furthermore, the DMN and the fronto-parietal network exhibited higher ICC coefficients than those in the BG and limbic networks (Figure S4C). Although Cohen’s *d* values reached zero when the TS method, but not other methods, was applied, the limited improvements of the ICC coefficients (6.1% to 7.0%) were also observed in rs-FCs. These results indicate that other confounds and state-like factors, such as arousal, might hinder the improvements of ICC coefficients in rs-FCs.

The current study confirmed the effectiveness of three harmonization methods on Cohen’s *d* values and classification accuracies. By applying the harmonization methods, Cohen’s *d* values were decreased in all the structural parameters and rs-FCs. For the TS and TS-Combat methods, classifiers could not distinguish participants from both protocols when data from 19 to 26 participants were available for estimating the protocol bias. We also observed the limited effectiveness of the Combat method, which is consistent with a prior study [Maikusa et al., 2021]. The limited efficacy might be attributed to the specificity of the dataset used in this study. Indeed, a previous study showed the effectiveness of the Combat method on rs-FCs similar to the TS method [Yamashita et al., 2019]. Further research is necessary to generalize our findings on other datasets for examining the upgrading effects.

The current study showed that the classification performance of SVM and ridge logistic regression decreased below the chance level when increasing the number of participants for estimating the effects, raising the possibility of information leakage due to overfitting. By plotting the distributions of posterior probabilities, we observed the flipped distributions of posterior probabilities, and this effect was more severe in rs-FCs with TS and TS-ComBat methods (Figures S1 and S2). Since the TS and TS-ComBat methods incorporate the participant factor into the GLM, the estimates of participant factor might re-introduce the protocol bias in the opposite direction, especially if the brain parameter with poor ICC coefficients (e.g., rs-FCs and CT) is used. Future research is necessary to investigate the cause of this phenomenon.

The current study showed the improved performance of age prediction, especially when applying LASSO with TS and TS-ComBat methods, except for rs-FCs and the Combat method. These observed improvements might be attributed to the increased consistency within the dataset, especially by applying TS and TS-ComBat methods. Indeed, SVR also showed limited improvements in the prediction performance. This might be due to the fact that SVR exploits the overall pattern as prediction while LASSO methods select the most reliable features within the dataset. In contrast to TS and TS-ComBat methods, the ComBat methods showed degradation of prediction performance on almost all brain parameters with the two prediction models, which are inconsistent with prior findings [Fortin et al., 2018; Yamashita et al., 2019].

The main reason for these observations might be attributed to the specificity of TS data used in this study and to the fact that the inclusion of age and sex as covariates of interest in the ComBat method might be not sufficient to characterize the individuals. It is important to investigate the generalizability of our findings on other datasets in future research.

The present findings have several limitations. First, we used two different scanning protocols with different head coils (i.e., 12-ch and 32-ch head coil). The difference in the number of channels affects the signal-to-noise ratio of images. We thus cannot rule out the possibility that, rather than protocol differences, differences in the head coils may hinder the improvements of test-retest reliability by applying the harmonization methods. Second, this study did not run MRI scanning twice with the same protocol to compute the test-retest reliability within the same protocol. We, thus, could not conclude that the harmonization methods completely eliminate the effects of protocol bias to achieve the level of true test-retest reliability. Third, we used the same preprocessing pipeline for both protocols instead of the state-of-art preprocessing pipeline [Glasser et al., 2013]. In addition, we did not apply ICA-FIX [Griffanti et al., 2014; Salimi-Khorshidi et al., 2014] or ICA-AROMA [Pruim et al., 2015a; Pruim et al., 2015b] to remove the effects of artefactual signals. A previous study demonstrated that ICA-based denoising could remove the site differences [Feis et al., 2015], suggesting that combining an ICA-based denoising method with a harmonization method might improve the test-retest reliability of rs-FCs. Further research is needed to investigate an optimal combination of preprocessing pipelines to mitigate protocol and scanner differences. Lastly, the current study did not examine the impact of protocol upgrades on other popular R-fMRI metrics, such as the amplitude of low-frequency fluctuations (ALFF) [Zang et al., 2007], fractional ALFF [Zou et al., 2008], degree centrality [Zuo et al., 2012], regional homogeneity [Zang et al., 2004], and voxel-mirrored homotopic connectivity [Zuo et al., 2010]. Further research is needed to systematically investigate the impact of protocol upgrades on other commonly-used R-fMRI metrics.

## Conclusion

We evaluated the effects of upgrading acquisition techniques on several brain parameters using univariate and multivariate analyses. Additionally, we showed the efficacy of three harmonization methods for mitigating the upgrading effects and the advantages of the TS and TS-ComBat methods over the ComBat method, at least in our dataset. The present findings provide implications for maintaining the continuity of MRI data before and after upgrading the MR system or acquisition techniques.

## Acknowledgments

This work was supported by the Japan Agency for Medical Research and Development (AMED; grant numbers: JP21dm0307008 to RH, JP19dm0307026 to DK and TI). This work was partially supported by the JSPS KAKENHI (19K03370 to TI, 21K15719 to YYA).

## Ethics approval statement

The study was approved by the Institutional Review Board of Showa University Karasuyama Hospital and was prepared in accordance with the ethical standards of the Declaration of Helsinki.

## Data availability

The datasets required the approval of an ethical review board. Please contact the corresponding author (T.I.).

