## Supplementary Information for "Effects of upgrading acquisition-techniques and harmonization methods: A multi-modal MRI study with implications for longitudinal designs"

**Title:**

**Short running title:**

Effects of protocol upgrades on MRI data

**Authors:**

Takashi Itahashi, Ph.D.[1], Yuta Y. Aoki, M.D., Ph.D.[1], Ayumu Yamashita, Ph.D.[2,3], Takafumi Soda, MSc.[4,5], Junya Fujino, M.D., Ph.D.[1,6], Haruhisa Ohta, M.D., Ph.D.[1], Ryuta Aoki, Ph.D.[1], Motoaki Nakamura, M.D., Ph.D.[1], Nobumasa Kato, M.D., Ph.D.[1], Saori C. Tanaka, Ph.D.[2], Daisuke Kokuryo, Ph.D.[7], Ryu-ichiro Hashimoto, Ph.D.[1,8]

**Affiliations:**

[1]Medical Institute of Developmental Disabilities Research, Showa University, Tokyo, Japan

[2]Brain Information Communication Research Laboratory Group, Advanced Telecommunications Research Institutes International, Kyoto, Japan

[3]Department of Psychiatry, Boston University School of Medicine, Massachusetts, USA

[4]Department of Information Medicine, National Institute of Neuroscience, National Center of Neurology and Psychiatry, Tokyo, Japan

[5]NCNP Brain Physiology and Pathology, Graduate School of Medical and Dental Sciences, Tokyo Medical and Dental University, Tokyo, Japan

[6]Department of Psychiatry and Behavioral Sciences, Graduate School of Medical and Dental Sciences, Tokyo Medical and Dental University, Tokyo, Japan

[7]Graduate School of System Informatics, Kobe University, Kobe, Japan

[8]Department of Language Sciences, Graduate School of Humanities, Tokyo Metropolitan University, Tokyo, Japan

**Correspondence:**

Takashi Itahashi, Ph.D.

Senior Assistant Professor

Medical Institute of Developmental Disabilities Research, Showa University, 6-11-11 Kita-karasuyama, Setagaya, Tokyo 157-8577, Japan.

**Pre-and post-upgrade reliability**

To assess the pre-and post-protocol-upgrade reliability, we computed intra-class correlation (ICC) coefficients, in which we treated pre- and post-upgrade measurements as raters. We observed negative ICCs, especially when computing ICCs for resting-state functional connections (rs-FCs). Because negative ICCs are difficult to interpret [Müller and Büttner, 1994; Rousson et al., 2002], prior test-retest studies have set negative ICCs to zero (i.e., completely non-reliable) [Braun et al., 2012; Kong et al., 2007]. In this study, we focused on the improvements of ICC coefficients rather than ICCs themselves. We, therefore, did not set negative ICCs to zero.

Based on previous studies [Cicchetti and Sparrow, 1981; Noble et al., 2019], we classified the ICC coefficients into four categories: poor < 0.4, fair 0.4 - 0.59, good 0.6 - 0.74, and excellent ≥ 0.75.

*Structural parameters:*

Figure S3 shows the results of ICC for CT, SA, and CV, respectively. The mean between-protocol reliability for CT was worst compared with other parameters (CT [mean ± SD]: 0.49 ± 0.18, range: -0.02 - 0.83; SA: 0.79 ± 0.15, range: 0.08 - 0.97; and CV: 0.75 ± 0.13, range: 0.23 - 0.96). Based on the ICC rating criteria (Figure S3D), CT exhibited fair ICC rating (poor: 29.75%, fair: 32.5%, good: 33.5%, and excellent: 4.25%), while the other parameters exhibited relatively better ICC rating (SA: poor: 2.5%, fair: 8.5%, good: 13.75%, and excellent: 75.25%; and CV: poor: 2.0%, fair: 9.75%, good: 33.25%, and excellent: 55%).

*rs-FCs:*

Figure S4A shows the results of ICC analysis for rs-FCs. The rs-FCs exhibited poor ICC coefficients than those of cortical parameters (rs-FCs [mean ± SD]: 0.31 ± 0.18, range: -0.42-0.87). Based on the ICC rating categories, rs-FCs exhibited dominantly poor ICC rating (poor: 69.08%, fair: 24.79%, good: 5.69%, and excellent: 0.44%) (Figure S4B). Within-network exhibited higher ICC coefficients compared to between-network ($p < 0.001$) (Figure S4C). In the within-network, default-mode (DMN) and fronto-parietal (FP) networks exhibited relatively higher ICC coefficients compared with other resting-state networks (RSNs), while the limbic and basal ganglia (BG) networks exhibited poor ICC coefficients.

**Effects of harmonization methods on ICCs**

*Structural parameters:*

We computed the ICCs after applying the harmonization methods (Figures S5A-C) to assess their effectiveness. All the harmonization methods improved the ICCs in all the structural parameters (CT: 45.2 ± 66.5% for TS and TS-ComBat methods, and 45.4 ± 63.6% for ComBat; SA: 13.4 ± 39.8% for TS, 13.3 ± 39.7% for TS-ComBat, and 12.3 ± 33.0% for ComBat; CV: 12.4 ± 18.6% for TS and TS-ComBat, and11.2 ± 16.4% for ComBat).

*rs-FCs:*

In contrast to the structural parameters, the improvements of ICC coefficients were limited even when increasing the number of participants used for estimating the protocol bias (TS: 6.2 ± 9.6 %; TS-ComBat: 6.1 ± 9.3%; and ComBat: 7.0 ± 9.4%) (Figure 5D).

**Table S1. The MRI scanning parameters used in this study.**

| | SRPBS | HARP |
|---|---|---|
| Head coil | 12-ch head coil | 32-ch head coil |
| | Functional MRI | |
| Total scan time [min:s] | 10:17 | 5:08 |
| # of runs | 1 | 2 runs for AP<br>2 runs for PA |
| Repetition time (TR)[ms] | 2,500 | 800 |
| Echo time (TE) [ms] | 30 | 34.4 |
| Multi-band factor | - | 6 |
| Field of view (FoV) [mm] | 212 | 206 |
| Matrix size | 64 × 64 | 86 × 86 |
| Flip angle [deg] | 80 | 52 |
| In-plane resolution [mm] | 3.3 × 3.3 | 2.4 × 2.4 |
| # of slices | 40 | 60 |
| Slice thickness [mm]# | 3.2 | 2.4 |
| Slice gap [mm] | 0.8 | 0 |
| Slice acquisition order | Ascending | Interleaved |
| Phase encoding direction | PA | AP/PA |
| Prescan normalize | Off | On |
| AutoAlign | - | Head > Brain |
| | Structural MRI | |
| Total scan time [min:s] | 9:50 | 5:22 |
| TR [ms] | 2,300 | 2,500 |
| TE [ms] | 2.98 | 2.18 |
| Inversion time [ms] | 900 | 1,000 |
| FoV [mm] | 256 | 256 |
| Base resolution | 256 | 320 |
| Flip angle [deg] | 9 | 8 |
| In-plane resolution [mm] | 1.0 × 1.0 | 0.8 × 0.8 |
| Slice thickness [mm] | 1.0 | 0.8 |
| # of slices | 240 | 224 |
| Prescan normalize | On | On |
| Distortion Correction | On (3D) | Off |
| AutoAlign | - | Head > Brain |
| PAT mode | None | GRAPPA |
| Accel. Factor PE | - | 2 |
| Ref. lines PE | - | 32 |

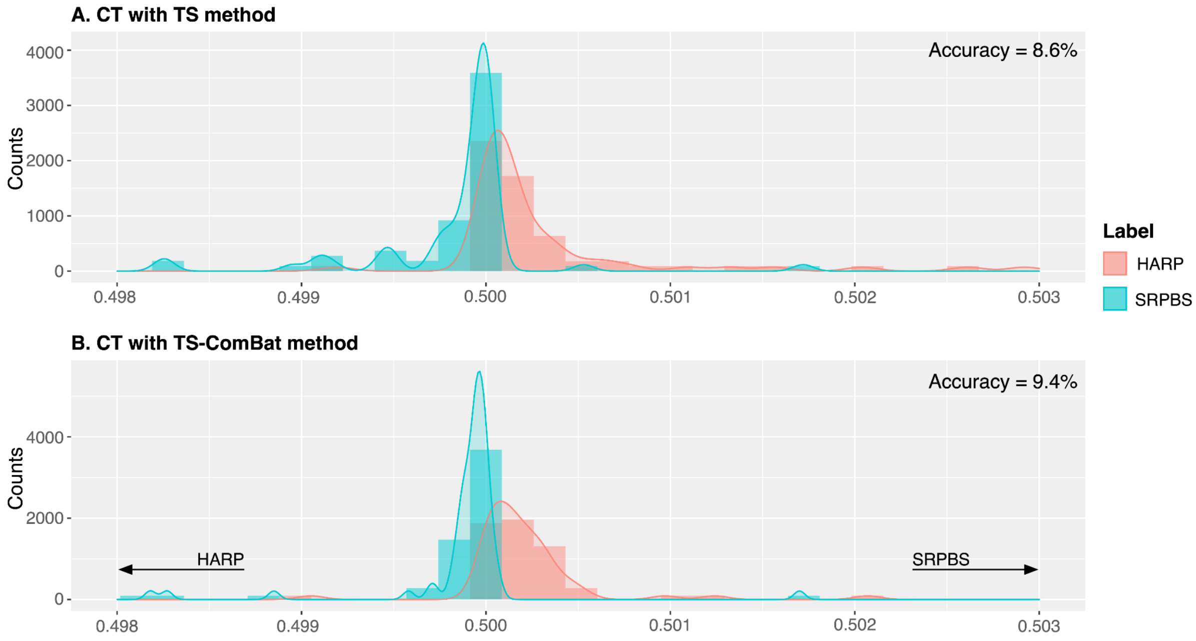


**Figure S1. The distributions of posterior probabilities obtained from ridge logistic regression with the cortical thickness (CT) after applying traveling subject (TS) and TS-ComBat methods.**

The upper panel (A) and lower panel (B) show the distributions of posterior probabilities after applying the TS and TS-ComBat methods, respectively. In both cases, the directions of the distributions of the posterior probability were reversed.

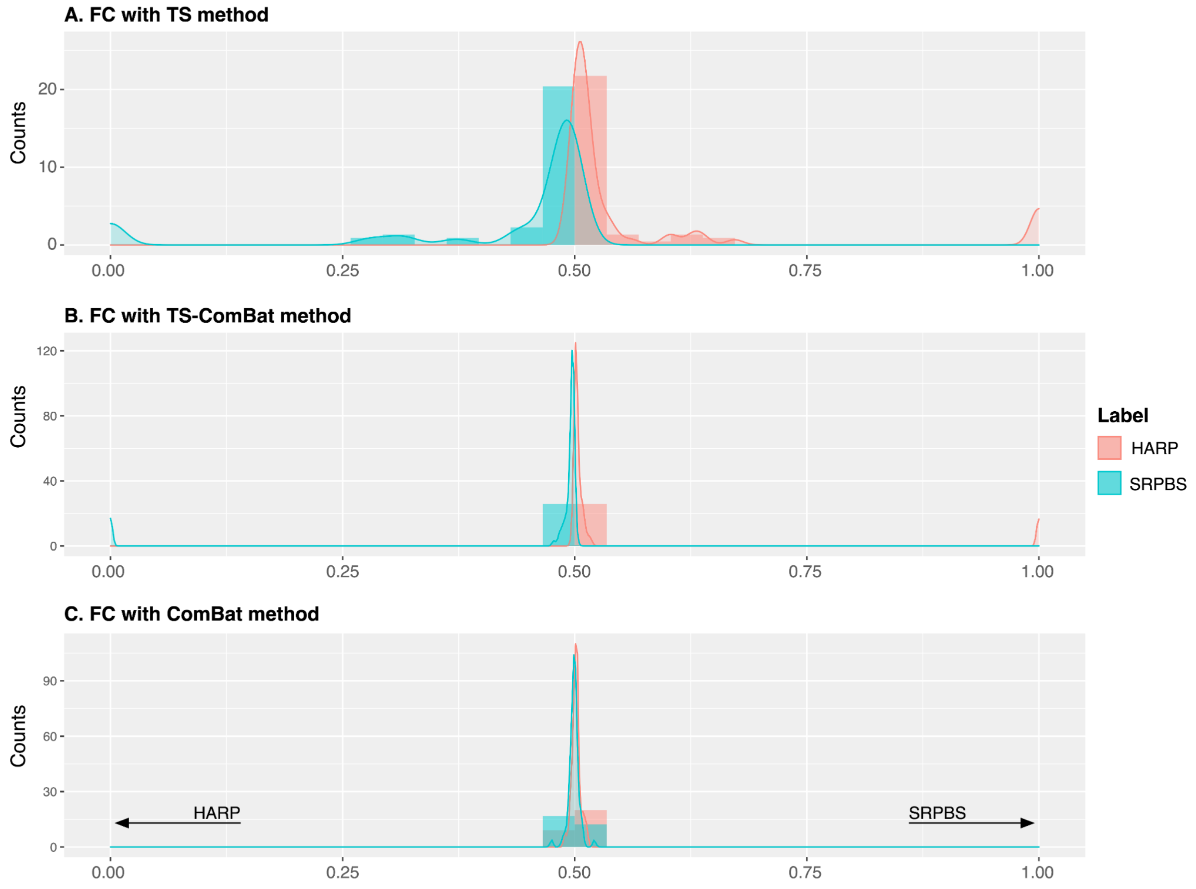


**Figure S2. The distributions of posterior probabilities obtained from ridge logistic regression with functional connectivity (FC) after applying harmonization methods.**

The upper panel (A) showed the distributions of posterior probabilities after applying the traveling subject (TS) method; the middle panel (B) showed the distributions of posterior probabilities after applying the TS-ComBat method; the lower panel (C) showed the distributions of posterior probabilities after applying the ComBat method. In both cases, the directions of the distributions of the posterior probability were reversed.

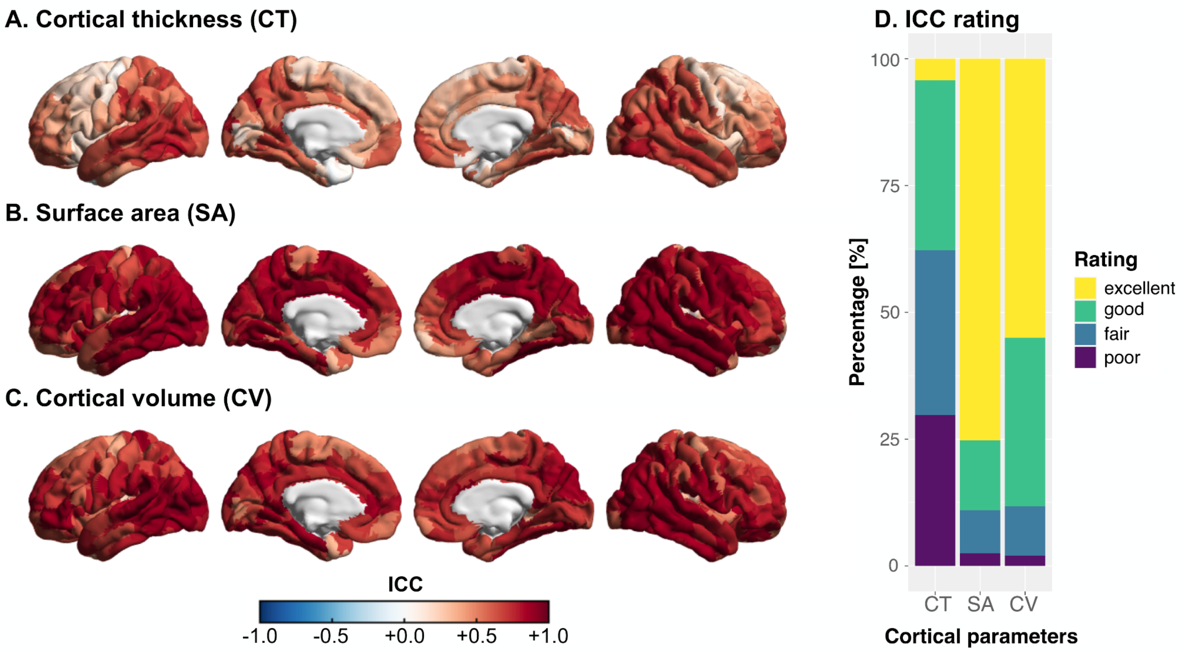


**Figure S3. The pre-and post-upgrade reliability on structural parameters.** (A) cortical thickness (CT), (B), surface area (SA), and (C) cortical volume (CV). The red and blue colors indicate the higher and lower ICC coefficients, respectively. (D) We classified The ICC coefficients into four categories: poor, fair, good, and excellent. SA and CV exhibited excellent reliability (SA: poor: 2.5%, fair: 8.5%, good: 13.75%, and excellent: 75.25%; CV: poor: 2.0%, fair: 9.75%, good: 33.25%, and excellent: 55%), while CT exhibited relatively fair reliability (poor: 29.75%, fair: 32.5%, good: 33.5%, and excellent: 4.25%).

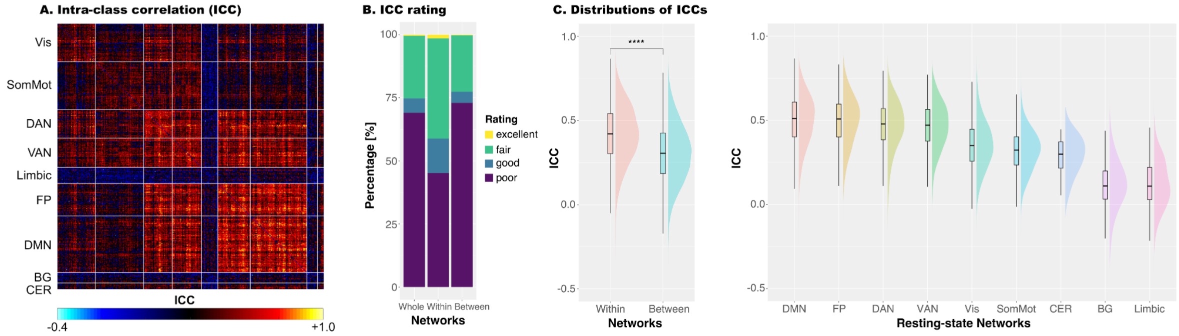


**Figure S4. The pre-and post-upgrade reliability on rs-FCs.** (A) The whole-brain pattern of Intra-class correlation coefficients (ICCs). (B) We classified The ICC coefficients into four categories: poor, fair, good, and excellent. (C) Comparisons of ICC distributions between within-network and between-network. Statistical comparisons exhibited statistically significant higher ICC coefficients in within-network compared to between-network. The right-most panel shows the distributions of ICC coefficients within each resting-state network (RSNs). Default mode (DMN) and fronto-parietal (FP) networks exhibited relatively higher ICC coefficients compared to other RSNs, while basal ganglia (BG) and limbic networks exhibited lower ICC coefficients. **Abbreviations:** BG: basal ganglia, CER: cerebellar, DAN: dorsal attention network, DMN: default mode network, FP: fronto-parietal network, SomMot: somatomotor network, VAN: ventral attention network, and Vis: visual network.

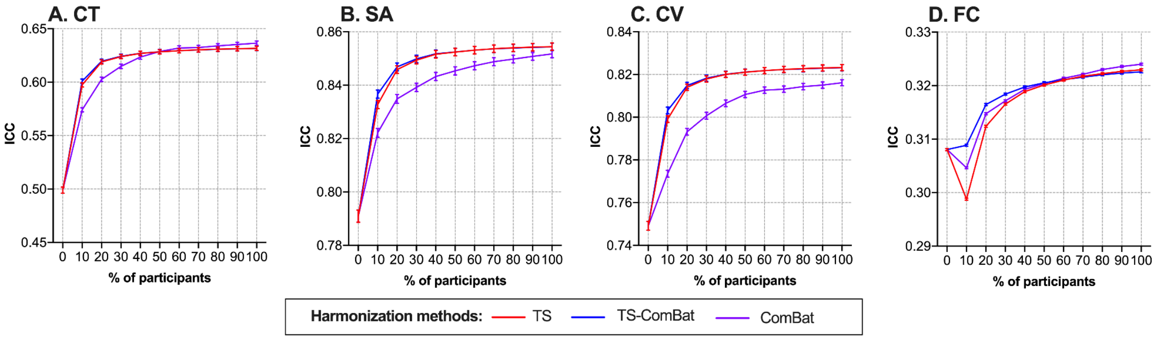


**Figure S5. The effects of harmonization methods on intra-class correlation (ICC) coefficients.**

The ICC coefficients were computed in the four brain parameters: (A) cortical thickness (CT), (B) surface area (SA), (C) cortical volume (CV), and (D) functional connectivity (FC). We varied the number of participants used for estimating the protocol bias from 0% to 100%. Of note, "0%" indicates that no harmonization methods are applied. The error bars showed the standard error of the mean (SEM). **Abbreviations:** TS: traveling subject.
